# Trajectory analysis of cardiovascular phenotypes from biobank data uncovers novel genetic associations

**DOI:** 10.1101/2020.05.10.087130

**Authors:** Tess D. Pottinger, Lorenzo L. Pesce, Anthony Gacita, Lindsey Montefiori, Nathan Hodge, Samuel Kearns, Isabella M. Salamone, Jennifer A. Pacheco, Laura J. Rasmussen-Torvik, Maureen E. Smith, Rex Chisholm, Marcelo A. Nobrega, Elizabeth M. McNally, Megan J. Puckelwartz

## Abstract

Approximately 6 million adults in the US have heart failure (HF). HF progression is variable due in part to differences in sex, age, and genetic ancestry. Previous population-based genetic studies have largely focused on cross-sectional data related to HF, a disease known to change over time. Utilizing longitudinal data trajectory probabilities as a continuous trait may increase the likelihood of finding significant, biologically relevant associations in a genome-wide association (GWA) analysis. We analyzed data from the electronic health record in a medical biobank from a single, metropolitan US center to gather clinically pertinent data for analyses. We evaluated whole genome sequencing of 896 unrelated biobank participants, including 494 with at least 1 electrocardiogram and 324 who had more than 1 echocardiogram (∼3 observations per person). A censored normal distribution multivariable mixture model was used to cluster phenotype measures for genome-wide analyses. GWA analysis on the trajectory probability of the corrected QT measurement (QTc) taken from electrocardiograms identified significant associations with variants in regulatory regions proximal to the *WLS* gene, which encodes the Wnt ligand secretion mediator, Wntless. *WLS* was previously associated with QT length using of approximately 16,000 participants supporting the utility of this method to uncover significant genetic associations in small datasets. GWA analysis on the trajectory probability of left ventricular diameter as taken from echocardiograms identified novel significant associations with variants in regulatory regions near *MYO10*, which encodes the unconventional Myosin-10. We found that trajectory probabilities improved the ability to discover significant and relevant genetic associations. This novel approach increased yield from smaller, well-phenotyped cohorts with longitudinal data from a medical biobank.

**AUTHOR SUMMARY:** Approximately 6 million adults in the US have heart failure, a disease known to change over time. In a hospital based electronic health record, electrocardiograms and echocardiograms, used to evaluate heart failure, can be tracked over time. We utilized these data to create a novel trait that can be applied to genetic analyses. We analyzed genome sequence of 896 biobank participants from diverse racial/ethnic backgrounds. Genome-wide association (GWA) analyses were performed on a subset of these individuals for heart failure outcomes. A statistical model that incorporates cardiac data that are tracked over time was used to cluster these data using a probabilistic approach. These probabilities were used for a GWA analysis for corrected QT measurement (QTc) and left ventricular diameter (LVID). The QTc interval analysis identified significant correlations with variants in regulatory regions near the *WLS* gene which encodes the Wnt ligand secretion mediator, Wntless. Analysis of LVID identified significant associations with variants in regulatory regions near the *MYO10* gene which encodes the unconventional Myosin-10. Through these analyses, we found that using the trajectory probabilities can facilitate the discovery of novel significant, biologically relevant associations. This method reduces the need for larger cohorts, and increases yield from smaller, well-phenotyped cohorts.

## INTRODUCTION

Genome-wide association studies (GWAS) have uncovered thousands of novel relationships between genetic variants and human disease conditions and traits. In order to achieve significance in the face of multiple testing correction, cohort sizes in the thousands or tens of thousands are generally required, presenting a challenge for rare disorders or for deeply phenotypically characterized cohorts. Because of the need for large sample sizes in GWAS, these studies generally often focus on case-control and cross-sectional data in curated cohort studies. A less commonly used approach in GWAS harnesses refined phenotyping that can be garnered from tracking traits over time, or longitudinal data analyses. Longitudinal data can be used to better identify patterns in complex traits, especially those with varying or late age of onset. Longitudinal data have not only been shown to increase power in GWAS but also have showed increases in heritability estimates in phenotypes such as blood pressure and body mass index (BMI) (Levy et al., 2000; Mei et al., 2012; Xu et al., 2014).

Medical biobanks provide a unique opportunity for longitudinal analyses. While medical biobanks are often not curated for a specific phenotype, they do benefit from having longitudinal medical record information. There are multiple approaches to analyzing longitudinal data. For example, longitudinal data can be clustered into distinct trajectories using a latent class mixed model (Proust-Lima et al., 2017). This technique was used to identify novel genetic variants associated with white blood cell count over time using a 2-group clustering on a cohort of approximately 10,000 individuals (Hall et al., 2019). An additional method used for clustering longitudinal data is a mixture model using latent class growth data (Jones and Nagin, 2007; Nagin, 2014). This method was applied in a 5-cluster model and used in a GWAS of blood pressure on a medium-sized dataset of approximately 1,000 individuals (Justice et al., 2016). This study utilized PROC TRAJ to capture the trajectory of the longitudinal data to be assigned to groups and relied on a censored normal distribution using multivariable mixture model (Jones and Nagin, 2007; Nagin, 2014). Clustering longitudinal data, while valuable, inherently loses data information by compressing longitudinal progression into groups. Alternative methods to evaluate longitudinal trajectories are needed improve robustness for even smaller datasets, making these smaller longitudinal datasets suitable for GWAS. The PROC TRAJ algorithm also provides probabilities that are used to assign an individual to a group. This probability can then be used to capture trajectory as a continuous trait when 2-group clustering is applied. We now utilized this method to analyze cardiac phenotypes from a medical biobank. Here we show that using trajectory probabilities as a continuous trait can improve the likelihood to find significant and biologically relevant associations for GWA analysis in small, longitudinal, well-phenotyped datasets.

Heart failure (HF) affects nearly 6 million adults (Benjamin et al., 2019; Mozaffarian et al., 2016). Within the U.S. population, there are differences in timing and progression of HF between individuals of different racial/ethnic groups and between males and females, and as the population ages the incidence of HF increases (Komanduri et al., 2017). The aging heart is characterized as having increased stiffness and changes in myocardial performance and size (Chiao and Rabinovitch, 2015; Oxenham and Sharpe, 2003). Electrocardiograms (EKGs) measure electrical conductance through the heart and yield significant continuous traits; progression of certain EKG measurements, such as the corrected QT interval (QTc) over time, serve as an indicator of pathology. The aging, diseased heart enlarges over time, and shifts in left ventricular internal dimensions (LVID) can be measured from echocardiograms.

Here, we used single nucleotide polymorphisms (SNPs) measured as part of whole genome sequencing on a diverse cohort of biobank participants from a single metropolitan site in the US. We used up to 23 years of longitudinal electrocardiogram and echocardiogram data from the electronic health record (EHR) to cluster participants into normal and atypical groups. We then utilized the trajectory probabilities that were estimate for these individuals to perform GWAS using ∼6.5 million SNPs. We found that variants near *WLS*, a gene which encodes Wntless, a Wnt ligand secretion mediator, were associated with QTc interval progression. We also found that variants in the *MYO10* gene, which encodes unconventional myosin-10, were associated with enlargement of the left ventricle over time. Integrated analyses of chromatin modifications and chromatin capture data suggested that SNPs near *WLS* and *MYO10* are likely regulatory in nature. Expression of both genes was increased in failed hearts compared to non-failed hearts. Notably, GWAS using only the clusters of normal and atypical values did not yield significant associations, while GWAS of trajectory probabilities did yield significant associations. These data suggest that trajectory probabilities of longitudinal data yield robust phenotypes that are likely to detect biologically relevant significant associations in GWA analyses.

## METHODS

### Cohort, clinical data and sequencing

NUgene is a biobank of adults who received care at Northwestern Medicine. The inclusion and exclusion criteria for enrollment in NUgene were previously described; primarily, this biobank recruits from the outpatient settings including but not limited to general medicine clinics (Ormond et al., 2009). Nine hundred NUgene biobank participants were selected to create a racially and ethnically diverse cohort, and DNA samples were whole genome sequenced (WGS), as described in (Pottinger et al., 2020). Variants were called using the Genome Analysis Tool Kit (GATK v3.3.0) best practices (Li and Durbin, 2010; McKenna et al., 2010). These analyses were conducted using the MegaSeq Pipeline (Puckelwartz et al., 2014). The NUgene Cohort files were anchored to the 1000 Genomes WGS data. Genetic population substructure was determined using the principal components calculated from the NUgene/1000 Genome data (**Supplemental Figure 1**). The first 3 principal components were used to capture the global genetic ancestry (PC1-PC3) of these individuals. Association analyses were conducted using PLINK v1.9 (Chang et al., 2015).

Echocardiogram, electrocardiogram, and demographic data were extracted from the electronic data warehouse (EDW) at Northwestern Medicine. Individual measures were obtained for left ventricular internal diameter-diastole (LVIDd) from echocardiogram reports that spanned as much as 14 years of echocardiogram data. The QTc interval was obtained from EKG measures and included multiple measures spanning as long as 23 years of data. Some participants had multiple measurements in a given year. For these participants the median value of results from that year was used for analysis. Calendar year was used as the time component for this analysis. Analyses were completed using SAS 9.4 and R v3.5.1. The diagnosis of heart failure was determined by ICD9 diagnosis codes 425 and all sub-codes, and ICD10 diagnostic codes I42 and all sub-codes.

### Trajectory Analysis

Trajectory analyses were conducted using a censored normal distribution multivariable mixture model on echocardiographic and electrographic quantitative traits using PROC TRAJ in SAS 9.4 (Jones and Nagin, 2007; Nagin, 2014). This method uses a likelihood function to assign a probability to a subject belonging to a trajectory cluster based on predetermined (k) number of clusters. This trajectory method is robust to missingness in longitudinal data. Consistent with the small sample size for this cohort, only 2 cluster separations were chosen. The model assigns a cluster to each individual as well as the probability of belonging in each cluster. Both measures (cluster and trajectory probability) were used in genome-wide associations.

### GWAS

Single nucleotide polymorphisms (SNPs) with a minor allele frequency of greater than 0.05 were used for GWA analyses. Variants were removed that had high deviations from Hardy Weinberg Equilibrium (threshold = 0.005) and genotype missingness rate greater than 0.05. Only autosomes were included for analysis. On average, this included 6.5 million SNPs per individual. All association analyses controlled for age, sex, and the first 3 principal components in an additive model. Quantile-quantile plots were generated for each association analysis to test for deviations from normalcy. Analyses were conducted using PLINK v1.9 (Chang et al., 2015). Sensitivity analyses were conducted to test for ancestry specific results.

### Variant fine-mapping

1000 Genomes data for all populations were used. A linkage disequilibrium (LD) map was created with the online tool (https://ldlink.nci.nih.gov/) using all individuals from the 1000 Genomes populations. Variants in LD with the top SNPs as well as those that were suggestively associated with outcome at the p-value<7E-07 threshold were further analyzed using the Roadmap epigenomics, ChromHMM, and ENCODE data repository to identify patterns indicative of enhancers and promoters using histone Chip-Seq data in heart left ventricle, aorta, and subcutaneous abdominal adipose tissues(2012; Kundaje et al., 2015). The “fold change over negative control” bigwig files were used from the histone H3K27Ac ChIP-Seq datasets. The files used for these analyses are described in Supplemental Table 1.

ATAC-Seq was conducted using iPSC cardiomyocytes. iPSCs were cultured and differentiated as described previously(Montefiori et al., 2018). ATAC-seq libraries were performed as in (Buenrostro et al., 2015) with minor changes as reported in (Montefiori et al., 2017). Briefly, 50,000 freshly harvested cardiomyocytes were pelleted by centrifugation, resuspended in 50 ul ice-cold lysis buffer (10mM Tris-HCl pH 7.4, 10mM NaCl, 3mM MgCl2, 0.1% Igepal CA-630) and centrifuged for 10 minutes at 4°C, 500 x g. Nuclei were resuspended in 50 ul of the transposase reaction buffer (Nextera DNA library prep kit (Illumina cat. FC-121- 1030) and 22.5 ul nuclease-free water) and incubated at 37°C for 30 minutes. DNA was purified from the reaction by purification on a Qiagen MinElute column (cat. 28206) according to manufacturer’s instructions and eluted in 10 ul of elution buffer (10mM Tris-HCl pH 8). Purified DNA was used immediately for PCR amplification using custom primers (Buenrostro et al., 2013). After 5 cycles, qPCR was used to determine the optimal number of additional PCR cycles required. Following PCR, ATAC-seq libraries were cleaned and size-selected using Ampure XP beads (Agencourt cat. A6388) to enrich for fragments between 300-700 bp and 1 ul was used for quantitation and size analysis on an Agilent Bioanalyzer High Sensitivity D1000 chip (Agilent cat. 5067-5584). ATAC-seq libraries were subjected to 50 bp single-end sequencing on an Illumina Hi-Seq 4000 instrument. A total of 8 technical replicates were generated from 3 different differentiation batches. All differentiations yielded CMs with purity > 80%. Reads were trimmed using cutadapt (Kechin et al., 2017) and aligned to the hg19 genome with Bowtie2 version 2.2.3 (Langmead and Salzberg, 2012) with default parameters. Reads with mapping quality lower than 10 were discarded. Mitochondrial reads and reads aligned to the same coordinates were removed. To call peaks, aligned reads from 8 technical replicates were pooled and input into MACS2 version 2.1.0 (Zhang et al., 2008) using the following parameters: -nomodel -llocal 20000 -shift -100 -extsize 200.

The CHiCAGO pipeline raw output of three replicates of iPSC-CM promoter capture Hi-C data were previously described (Cairns et al., 2016; Montefiori et al., 2018). Promoter-baited interacting fragments were filtered and 1kb was added to both ends of regions interacting with gene promoters due to the use of a 4-bp cutter (Mbol) to generate the Hi-C libraries. This method generates fragments that are on average 400bp, making it likely that neighboring fragments would contribute to the same interaction. Therefore, each interacting end was extended by 1kb to avoid over filtering while obtaining reproducible interactions between replicates. Data from each replicate was evaluated using the program Bedtools (Quinlan and Hall, 2010), and genomic interactions that were present in at least two replicates were retained. Bed files representing Hi-C interactions were visualized in the UCSC genome browser. The files used for these analyses are described in Supplemental Table 1. The VISTA Browser was used to test for H3K27ac patterns in fetal heart tissue, heart tissue of individuals with dilated cardiomyopathy, and tissue from normal hearts (Spurrell et al., 2019). The UCSC human genome browser was used to display the sequencing peaks.

### Transcription Factor Binding Sites and eQTL

Variants were then analyzed using JASPAR’s analysis tool for transcription factor binding sites in humans. A 30bp region around the reference and alternative variants in these regions were evaluated. Reference regions that showed differences in transcription binding in comparison to the alternative regions were further evaluated (Fornes et al., 2020).

Additionally, the GTEx eQTL calculator was used to evaluate the variants of interest in *WLS* and *MYO10* and the tissue information available for heart atrial appendage, heart left ventricle, skeletal muscle and artery (obtained from the GTEx Portal on 3/31/2020). Gene expression data from the GTEx browser was also used to evaluate the expression of these genes in a subset of tissues.

### RNA-Seq for Gene Expression Analysis in Heart Tissue

RNA-Seq analyses were conducted using samples from healthy and failed left ventricle obtained from failed transplants or as discarded tissue. These analyses have been previously described (Gacita et al., 2020). Genes with <1 count per million were removed from the analysis. Differentially expressed genes were defined as any gene with an FDR-corrected p-value of < 0.05.

## RESULTS

### Biobank-derived genomic and clinical findings

The NUgene biobank was established in 2000 and the electronic data warehouse aggregates clinical data from 1996. We used WGS data from 895 biobank participants; the demographics of this population was previously described (Pottinger et al., 2020). Using genetic ancestry, we found that approximately 46% of participants in the cohort used for genome-wide association analyses were of African ancestry (**Table 1**).

**Table 1.**
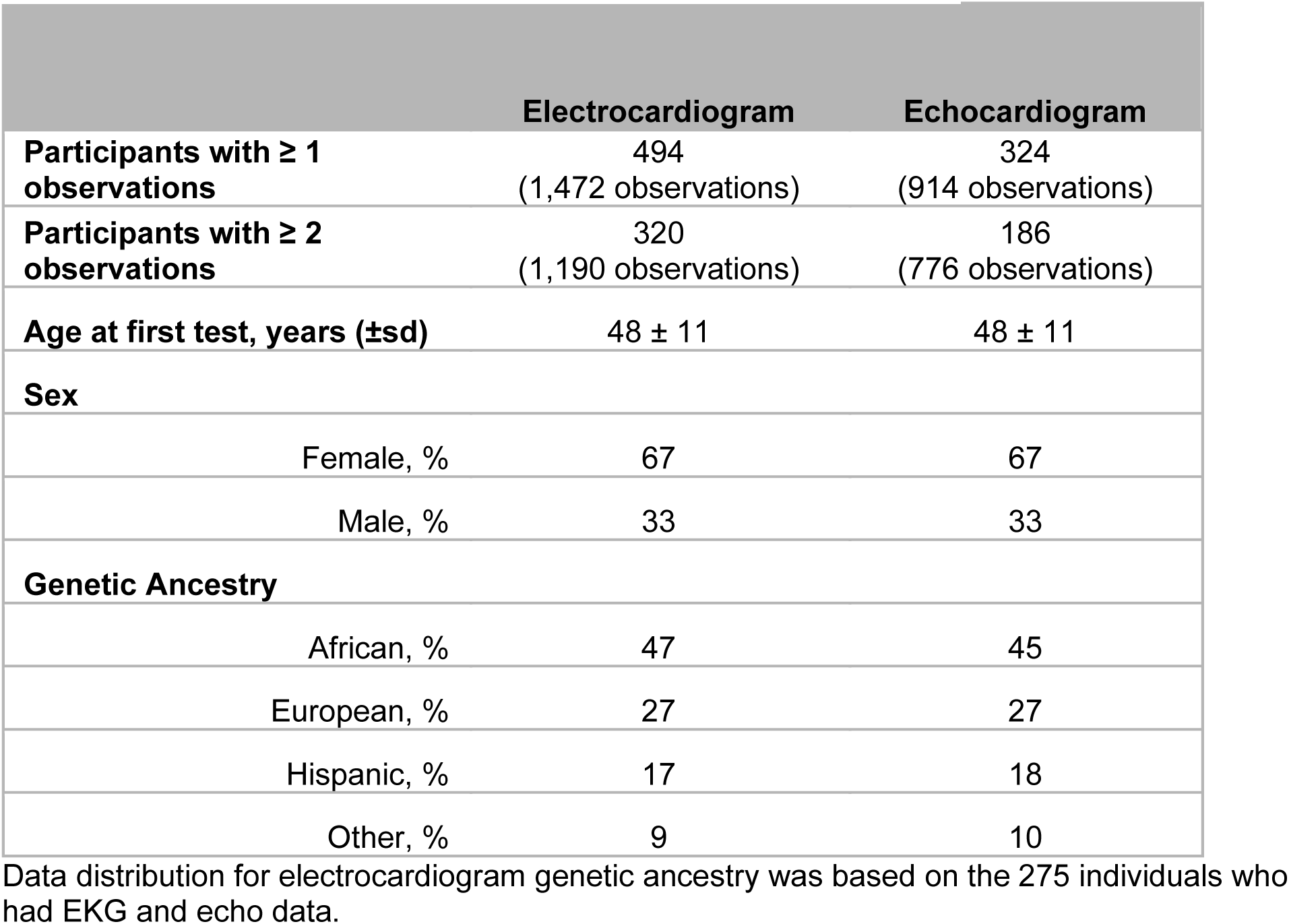
Demographic characteristics of the NUgene Cohort.

|  | **Electrocardiogram** | **Echocardiogram** |
| --- | --- | --- |
| **Participants with $\geq 1$ observations** | 494<br>(1,472 observations) | 324<br>(914 observations) |
| **Participants with $\geq 2$ observations** | 320<br>(1,190 observations) | 186<br>(776 observations) |
| **Age at first test, years (±sd)** | 48 ± 11 | 48 ± 11 |
| **Sex** |  |  |
| Female, % | 67 | 67 |
| Male, % | 33 | 33 |
| **Genetic Ancestry** |  |  |
| African, % | 47 | 45 |
| European, % | 27 | 27 |
| Hispanic, % | 17 | 18 |
| Other, % | 9 | 10 |

Data distribution for electrocardiogram genetic ancestry was based on the 275 individuals who had EKG and echo data.

Of the 895 participants with WGS data, 494 had an EKG with the QTc interval measured. Over the 23 years of data collection, these 494 had an average of four QTc observations, with 320 participants having at least two measures of QTc (**Table 1**). Three hundred and twenty-four individuals had echocardiogram data that included a measurement of the left ventricle internal dimension in diastole (LVIDd). All LVIDd measures were normalized to body surface area (BSA). On average, the 324 participants had 4 independent measures of LVIDd over a maximum of 14 years of information. Of these individuals, 186 participants had at least 2 measures of LVIDd. LVIDd is a standard measure of left ventricular size. The average age for the first electrocardiogram and echocardiogram test in this cohort was 48 years; and 67% of participants were female (**Table 1**).

### Trajectory probabilities of QTc interval identify novel variants in GWAS

The longitudinal data from the EHR were clustered in two groups: a normal QTc interval cluster, with data that ranges from 420 to 440 ms, and long QTc interval cluster, with data that ranges from 460 to 540 ms. The majority of participants clustered into the normal QTc interval cluster (**Figure 1A**), while 16% (44 of 275 subjects) were assigned to the long QTc interval cluster. Figure 1B shows the probability density plot for these same values, revealing the majority of participants fell into the ‘normal’ cluster (physiologically normal range of values) with a considerable number of individuals falling between the two tails (**Figure 1B**). We analyzed only individuals with both an EKG and echocardiogram, as this may indicate a higher likelihood or suspicion of cardiac disease. Of the 275 subjects in this subset, 89 had a diagnosis of heart failure. Of these 89, 29 were included in the long QTc cluster and 60 were contained in the normal cluster. The cluster assignment and probability density showed similar distribution between these individuals and the original 494 individuals.

**Figure 1.**
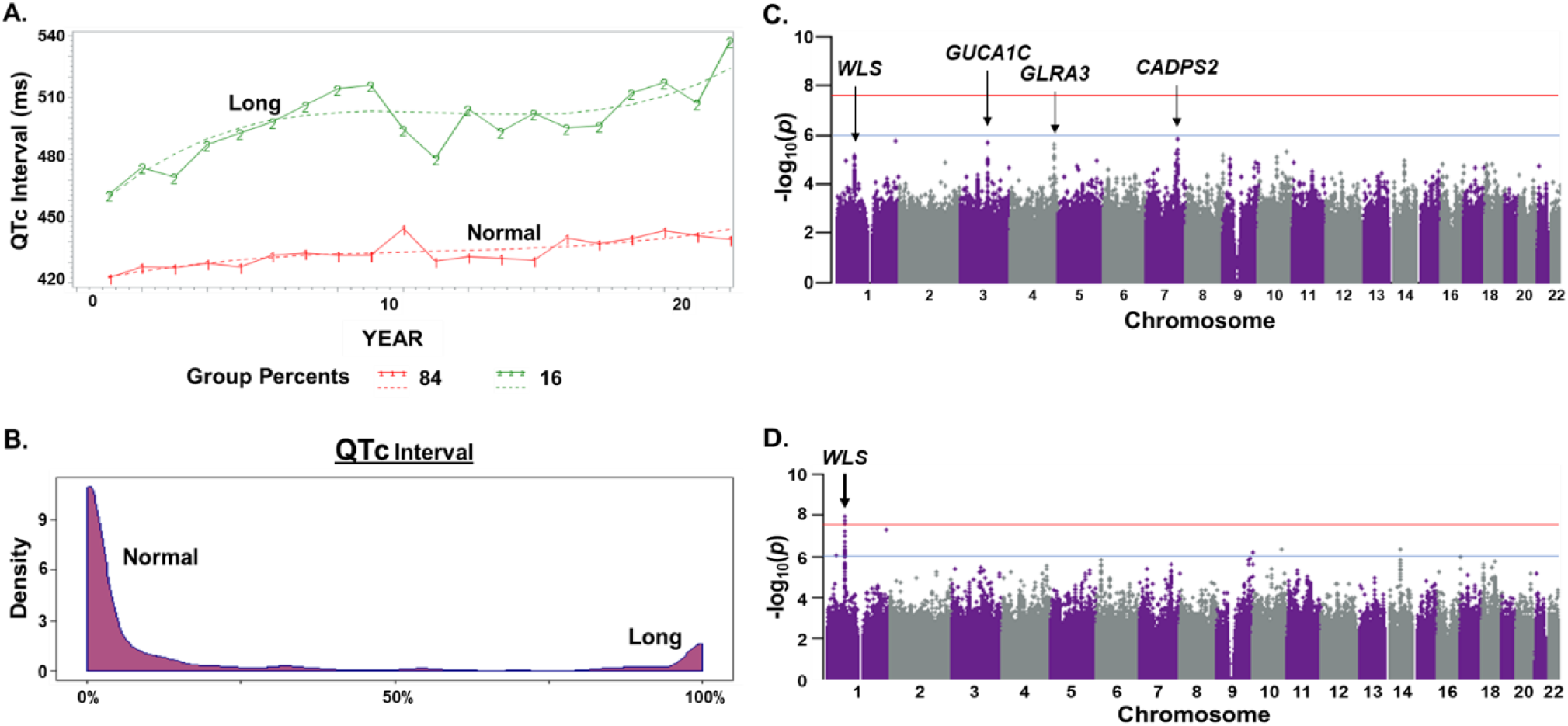
Continuous, but not binary, analysis of QTc intervals identifies genome-wide significance at *WLS*. **A.** A binary analysis of QTc trajectory was derived from EKGs over 20 years (N=275 subjects). This data was clustered into long and normal QTc intervals using a censored normal distribution using multivariable mixture model. With this clustering, group 1- normal (red line) included 84% of the cohort and, group 2-long (green line) included 16% of the cohort. **B.** Probability estimates, distributed between 0 and 100%, were used to distribute measures of QTc interval into 2 groups, normal and long QTc. The probability of clustering into these 2 groups showed a bimodal distribution, with a small number falling between the peaks. Using all probability estimates, including those between the two tails, created a trajectory probability. **C)** GWAS of QTc was conducted using the binary data in **A**. This analysis controlled for age, sex, and global genetic ancestry (PC1-PC3). The red line indicates genome-wide significance (p-value<3E-08); the blue line represents the suggestive line for significance (p-value<7E-07). Non-significant association was seen for variants in or near *WLS, GUCA1C, GLRA3*, and *CADPS2.* **D)** In contrast, GWAS of QTc interval trajectory probability shown in B was performed with the same subjects and identified a significant association with variants in or near *WLS.* The analysis was similarly controlled for age, sex and global genetic ancestry (PC1- PC3) as in C, and the same genome-wide significance is marked by the red and blue lines. *WLS* encodes the Wnt ligand secretion mediator, Wntless.

We conducted a GWAS using the QTc cluster data from Figure 1A on the 275 individuals with EKG and echocardiogram data. All GWAS controlled for age, sex, and global genetic ancestry (PC1-PC3) (**Figure 1C**), although not differences were seen in trait distribution by sex (**Supplemental Figure 2**). This approach, using the 2-cluster phenotyping, yielded only non-significant associations at a p-value<3E-08 threshold in and around *WLS, GUCA1C, GLRA3*, and *CADPS2*. This analysis showed nominal deviations from normal with a lambda of 1.04 (**Supplemental Figure 3A**) as a lambda between 1 and 1.05 is considered the acceptable threshold for deviation (Freedman et al., 2004). In contrast, a GWA using trajectory probabilities to calculate QTc distribution as a continuous trait showed a significant association (p-value<3E-08) with SNPs near the *WLS* gene (**Figure 1D**). This analysis showed minimal deviations from normal with a lambda of 1.01, an improvement from the dichotomous analysis (**Supplemental Figure 3B**). These analyses indicate that trajectory probability outcomes can be used to detect novel significant associations of longitudinally well-phenotyped cohorts.

The variants that were significantly associated with QTc trajectory probability and those that were suggestively associated (p-value<7E-07) were further investigated. All risk alleles of these variants show an increase in QTc interval probability, or likelihood of having a long QTc interval, in individuals with these variants (**Table 2**). The risk allele frequency for these variants range from 0.2 to 0.4. A similar direction of association is observed when evaluating these variants in the dichotomous or cluster association analysis indicating that the risk allele is associated with a longer QTc interval (**Supplemental Table 2)**. To further investigate these results, a variant-specific association analysis was performed evaluating the QTc interval value cross-sectionally using the first EKG value. These data also show a similar direction of association as the trajectory probability analysis emphasizing the robustness of the associations. Ancestry-specific analyses also showed similar trends in association for these variants (**Supplemental Table 3**) supporting the value of this method.

**Table 2.** Characteristics of variants in the *WLS* peak region associated with QTc interval trajectory probability

| **Variant** | **rs number** | **Risk Allele** | **Parameter estimate** | **P-value** | **Nearest gene** | **Distance from TSS** | **Global AF** |
| --- | --- | --- | --- | --- | --- | --- | --- |
| **chr1:68762502** | rs1304851468 | G | 0.2 | 1.1E-8 | *WLS* | 64KB | 0.4* |
| **chr1:68788564** | rs4536035 | C | 0.19 | 1.83E-8 | *WLS* | 90KB | 0.2 |
| **chr1:68788712** | rs4506498 | T | 0.19 | 2.52E-8 | *WLS* | 90KB | 0.3 |
| **chr1:68785213** | rs11209261 | T | 0.18 | 6.31E-8 | *WLS* | 87KB | 0.3 |
| **chr1:68764949** | rs10889761 | T | 0.18 | 10E-8 | *WLS* | 67KB | 0.3 |
| **chr1:68797643** | rs10789260 | G | 0.19 | 1.85E-7 | *WLS* | 99KB | 0.2 |
| **chr1:68804349** | rs4926357 | G | 0.18 | 3E-7 | *WLS* | 106KB | 0.2 |
| **chr1:68787424** | rs11309689 | AT | 0.15 | 4.24E-7 | *WLS* | 89KB | 0.3 |
| **chr1:68784444** | rs6588315 | G | 0.17 | 4.32E-7 | *WLS* | 86KB | 0.3 |
| **chr1:68804055** | rs4926330 | A | 0.18 | 5.26E-7 | *WLS* | 106KB | 0.3 |
| **chr1:68794008** | rs4598529 | T | 0.18 | 5.53E-7 | *WLS* | 96KB | 0.2 |
| **chr1:68787891** | rs4244620 | C | 0.16 | 6.34E-7 | *WLS* | 90KB | 0.3 |
| **chr1:68804530** | rs4926360 | T | 0.18 | 6.95E-7 | *WLS* | 106KB | 0.2 |
| **chr1:68804258** | rs4926356 | A | 0.18 | 8.08E-7 | *WLS* | 106KB | 0.2 |

TSS: transcription start site; AF: Allele frequency from gnomAD
\*Based on rs199703974, rs1304851468 absent from gnomAD data

### Enhancers of *WLS* are associated with cardiac outcomes

Multiple SNPs in an intergenic region on chromosome 1 near *WLS*, a gene encoding Wntless, a Wnt receptor that participates in Wnt secretion and recycling (Port and Basler, 2010) were associated with QTc interval. **Figure 2** shows the linkage disequilibrium (LD) map of the significantly associated QTc SNPs (p-value<3E-08) including rs199703974, rs4536035, rs4506498. Variants rs4536035 and rs4506498 form an LD block with variant rs4244620 which showed suggestive significance in the GWAS (p-value<7E-07).

**Figure 2.**
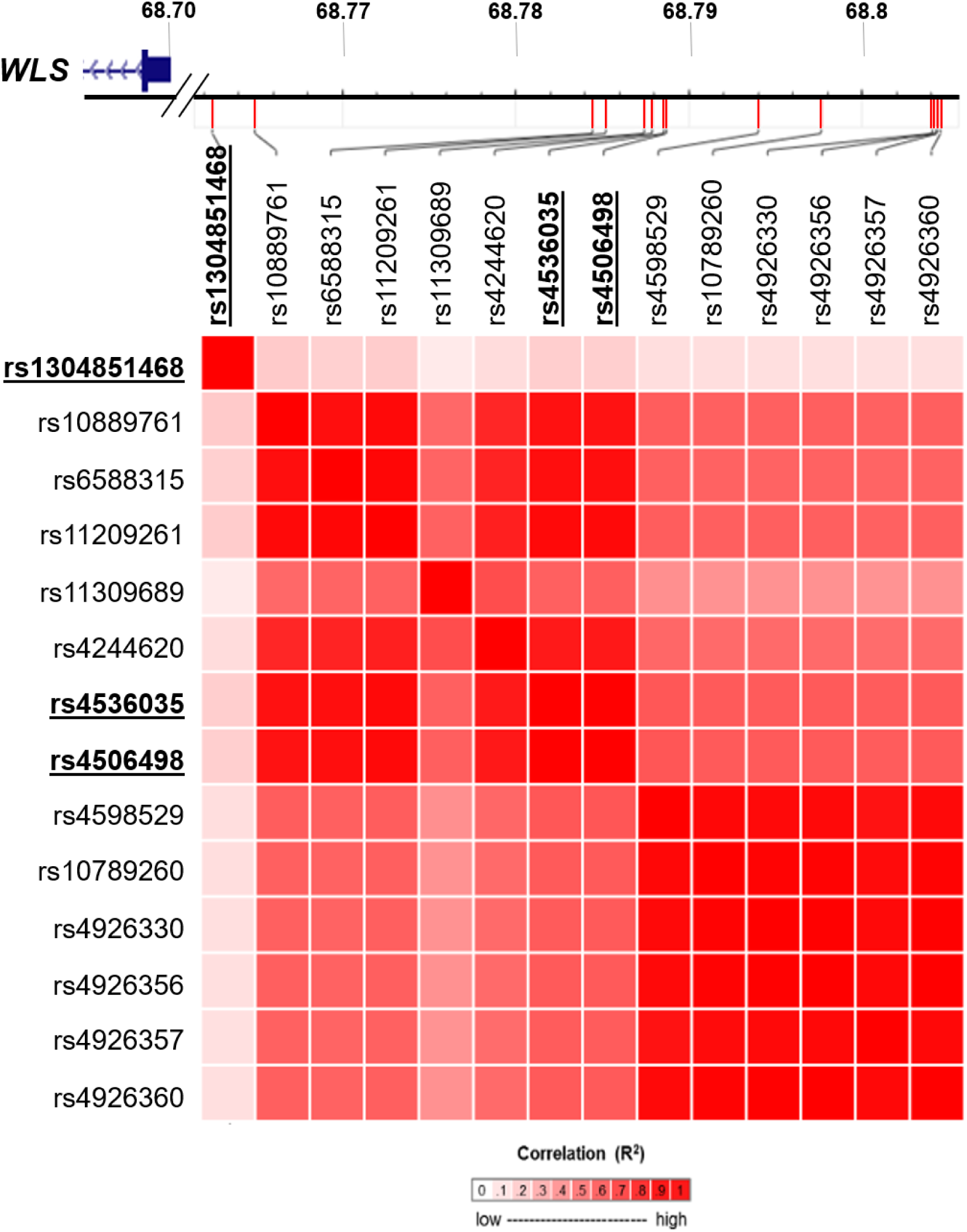
Linkage disequilibrium (LD) structure and location of SNPs associated with QTc trajectory probability near *WLS*. The position of WLS is shown at the top. Shown are the SNPs on chromosome 1 in LD with SNPs associated with QTc interval trajectory probability in all 1000 Genomes Populations (https://ldlink.nci.nih.gov/). The SNPs that were significantly associated (p-value<3E-08) with QTc interval trajectory probability; rs199703974, rs4536035, rs4506498; are bolded and underlined. Variants rs4536035, rs4506498 form an LD block with variant rs4244620 which showed suggestive significance in the GWAS (p-value<7E-07).

These variants map upstream of *WLS*, with the closest SNP 64KB 5’ from the *WLS* transcriptional start site (**Table 2**). ChIP-seq for H3K27Ac, a chromatin marker for enhancers, from human heart left ventricle revealed enrichment for H3K27Ac in and around variants rs1304851468 (**Figure 3A**, blue track, yellow box). As *WLS* is expressed broadly, and cardiovascular disease derives from both cardiac intrinsic and extrinsic factors, we also evaluated H3K27Ac data from human aorta and subcutaneous abdominal adipose tissue. ATAC-seq, which assesses open chromatin, was evaluated from cardiomyocytes differentiated from induced pluripotent stem cells (iPSC-derived cardiomyocytes). This analysis also supported the presence of potential active enhancer peaks in and around variants rs1304851468 and rs4244620 (**Figure 3A**, red track). Chromatin capture – promoter interaction data was similarly queried. We also evaluated the pattern of long-range promotor-enhancer interactions in iPSC-derived cardiomyocytes (Montefiori et al., 2018). These data showed looping of the region near rs4244620 to the *WLS* promotor, further supporting the notion that this region corresponds to a *WLS* cardiac enhancer (**Figure 3A**, purple track). Interactions were not observed for rs1304851468 and the *WLS* promotor. This may be due to a lack of interaction between the loci, or technical aspects such as the immaturity of the iPSC-derived cardiomyocyte model or an unresolvable distance based on the limits of the methods. Similar patterns of H3K27Ac acetylation were seen in both normal hearts and failed hearts (**Supplemental Figure 4**) (Spurrell et al., 2019). These analyses suggest that SNPs identified by GWAS map to potential *WLS* regulatory regions.

**Figure 3.**
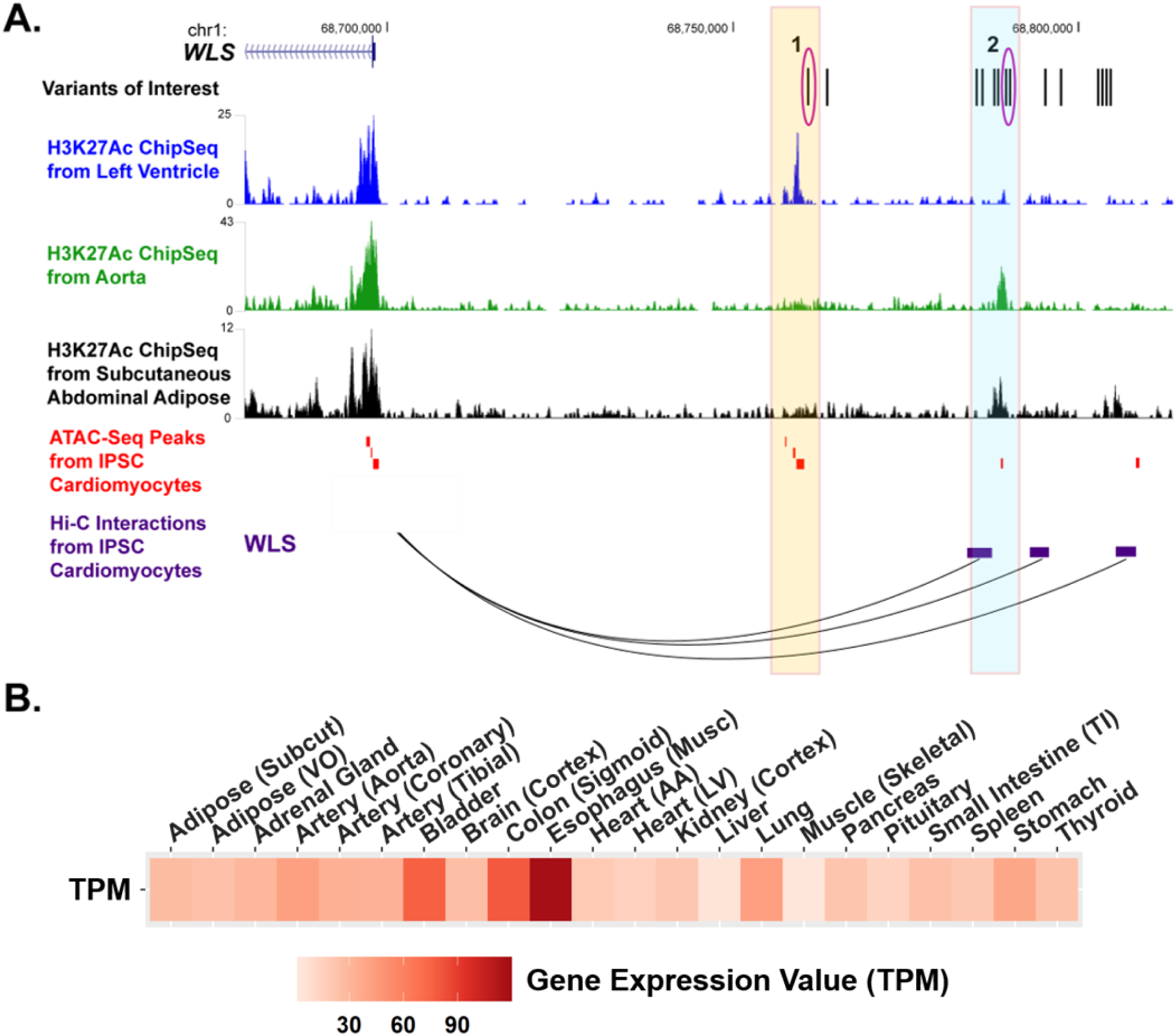
QTc-associated SNPs map to candidate *WLS* enhancer regions. **A.** QTc-associated SNPs are denoted by black lines as variants of interest. Significant SNPs are circled in magenta. The yellow and light blue boxes encompass sequences with molecular markings consistent with active enhancer elements. H3K27Ac ChIP-Seq data from left ventricle of human heart (bright blue track) showed enrichment in and around variants rs1304851468, indicated by magenta circle 1. Similarly, H3K27Ac ChIP-Seq data from human aorta (green track) and subcutaneous abdominal adipose tissue (black track) show enrichment in and around rs4244620, a variant in high LD with variants significantly associated with QTc interval trajectory (magenta circle 2). ATAC-seq from iPSC-derived cardiomyocytes (red track) enrichment in and around variants rs1304851468 and rs4244620. Chromatin capture of this region in iPSC-derived cardiomyocytes revealed a putative enhancer region for *WLS* near rs4244620 (purple boxes). H3K27Ac data from the Roadmap Epigenome Consortium. **B**. GTEx median expression in transcripts per million (TPM) of *WLS* across all available tissue types (data from the GTEx Portal on 3/28/2020). Abbreviations: Subcut-subcutaneous; VO-visceral(omentum); Musc-muscularis; AA-atrial appendage; LV-left ventricle; TI-terminal ileum

We compared WLS expression in failed hearts (n=97) and normal hearts (n=108) in a previously described cohort (Heinig et al., 2017). *WLS* was modestly but significantly differentially expressed between failed hearts and non-failed hearts (log fold change=0.078, Benjamini–Hochberg adjusted p=1E-8). The *WLS* gene is ubiquitously expressed with high expression in esophagus, bladder and aorta (**Figure 3B**). An eQTL analysis using GTEx data showed a significant association with *WLS* and rs4536035, a variant significantly associated with QTc interval trajectory probability, in the coronary artery (p-value=0.024). These analyses indicate that *WLS* may act through a vascularly driven process to affect cardiovascular outcomes.

### Trajectory probabilities of left ventricular size identify novel variants in GWAS

We tested for association with echocardiographic measures using longitudinal LVIDd data (corrected for BSA). The longitudinal data was grouped into 2 clusters that divided participants into a normal LVIDd cluster (ranging from 1.9 to 2.3 cm/m^2^) and dilated LVIDd cluster (ranging from 2.6 to 2.9 cm/m^2^). The normal cluster included 237 of 324 participants (73%) (**Figure 4A**), while 87 of 324 (27%) were assigned to the dilated cluster. A probability density plot for these values demonstrated a bimodal distribution with many individuals falling between the peaks (**Figure 4B**). From the total group, 89 had a diagnosis of heart failure with 35 in the dilated cluster and 56 in the normal cluster.

**Figure 4.**
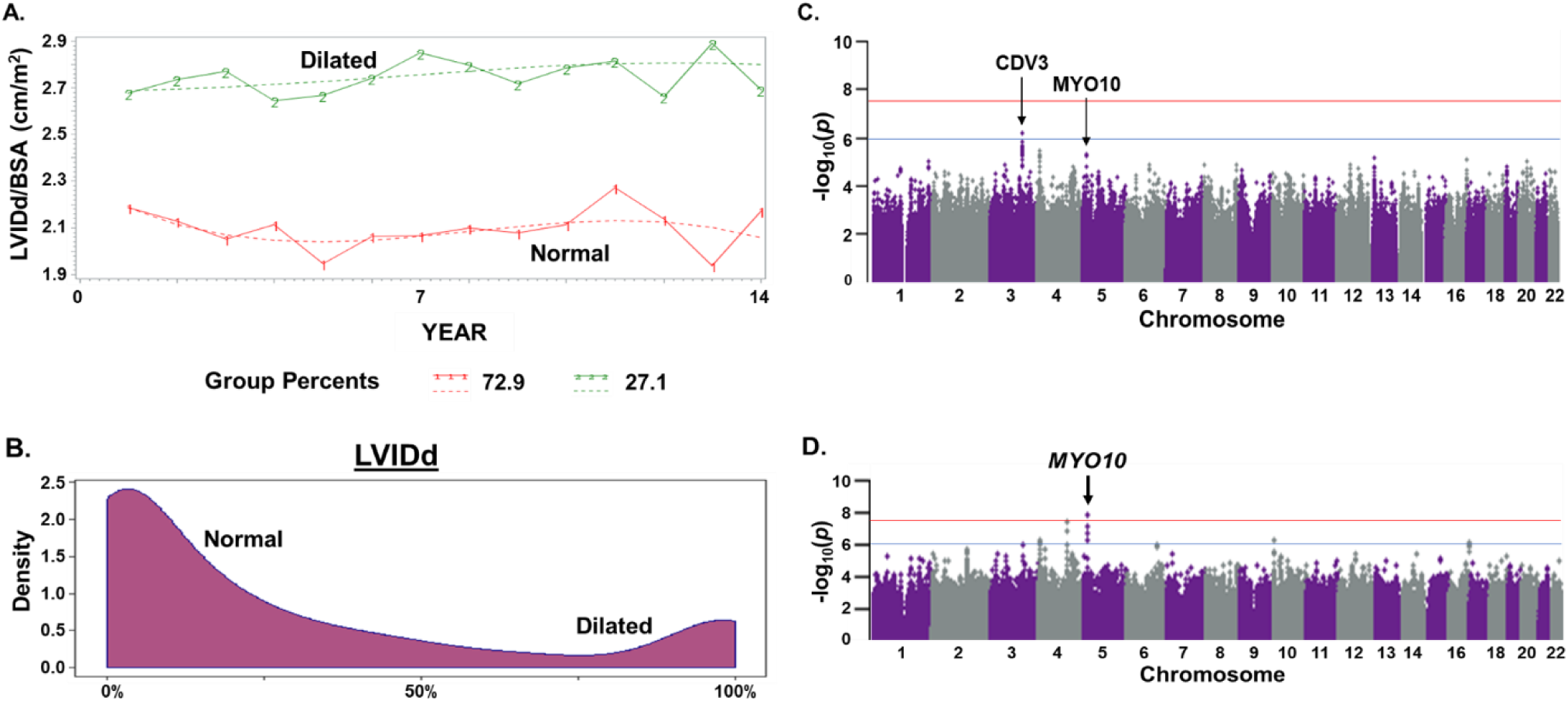
Continuous, but not binary analysis, of left ventricular dimensions identifies an association with *MYO10*. Left ventricular internal dimension in diastole (LVIDd) was extracted from echocardiogram reports over a 14 year interval (N=324 subjects). For all analyses, only LVIDd normalized to BSA was used. **A.** LVIDd measures were clustered into dilated and normal clusters using a censored normal distribution using multivariable mixture model. With this approach, 73% were in the normal (red line) group and 23% were in the dilated group (green line). **B.** The same data was analyzed by probability estimates distributed between 0 and 100%. The probability of clustering into normal and dilated showed a bimodal distribution, with a range of data falling between the two peaks. Using all probability estimates, including those between the two tails, created a trajectory probability that was used for GWAS. **C.** GWAS of LVIDd using the data in **A** showed a non-significant association with variants in or near *MYO10*, and *CDV3*. Analysis controlled for age, sex, and global genetic ancestry (PC1-PC3). **D.** GWAS of LVIDd trajectory probability controlling for age, sex, and global genetic ancestry (PC1-PC3) was performed with data in **B**. Significant associations were identified with variants in or near *MYO10;* which encodes myosin 10, a member of the myosin superfamily. The red line indicates the significance threshold line (p-value<3E-08); blue line represents the suggestive line for significance (p-value<7E-07).

A GWA analysis using 324 individuals with echocardiographic measures (**Table 1**) was conducted using the LVIDd clusters in Figure 4A controlling for age, sex, and global genetic ancestry (PC1-PC3), although not differences were seen in trait distribution by sex (**Supplemental Figure 2**). No significant associations were detected. However, suggestive association peaks were seen in and around *CDV3* and *MYO10*. This analysis showed nominal deviations from normal with a lambda of 1.03 (**Supplemental Figure 5A**). In contrast, using the trajectory probabilities from Figure 4B as a quantitative trait, a GWA analysis showed a significant association (p-value<3E-08) overlapping the *MYO10* peak from the dichotomous GWA analysis (**Figure 4D**). This analysis showed no deviation from normal with a lambda of 1 (**Supplemental Figure 5B**). These analyses again support that trajectory probability outcomes provide a novel method for which to find significant associations in GWAS of longitudinally well-phenotyped cohorts. All GWA were controlled for age, sex, and global genetic ancestry (PC1- PC3).

*MYO10* encodes myosin 10, a member of the myosin superfamily and has not previously been associated with echocardiographic measures. The risk alleles of the SNPs in this region associated with LVIDd probability, or higher likelihood of having a dilated LVIDd, are shown in **Table 3**. The minor allele frequency for these variants range from 0.05 to 0.07. A similar direction of association is observed when evaluating these variants in the dichotomous or cluster association analysis indicating that the risk allele is associated with a dilated LVIDd (**Supplemental Table 4)**. To further investigate these SNPs, a variant-specific association analysis was used evaluating the LVIDd value cross-sectionally using the first echocardiogram value. These data also show a similar direction of association as the trajectory probability analysis. These analyses indicate that this method is robust in finding significant associations in GWA analyses. Ancestry-specific analyses also showed similar trends in association for these variants (**Supplemental Table 5**) supporting the utility of this method.

**Table 3.** Characteristics of variants in the *MYO10* peak region associated with LVIDd trajectory probability

| Variant | rs number | Risk Allele | Parameter estimate | P-value | Nearest gene | Distance from TSS | Global AF |
| --- | --- | --- | --- | --- | --- | --- | --- |
| **chr5:16957078** | rs113861938 | T | 0.42 | 2.10E-9 | *MYO10* | 21KB | 0.05 |
| **chr5:16956160** | rs112072727 | T | 0.33 | 1.27E-8 | *MYO10* | 20KB | 0.06 |
| **chr5:16956338** | rs78565023 | G | 0.32 | 1.41E-8 | *MYO10* | 20KB | 0.06 |
| **chr5:16956916** | rs77050123 | A | 0.31 | 6.49E-8 | *MYO10* | 20KB | 0.06 |
| **chr5:16958033** | rs7734901 | G | 0.3 | 7.46E-8 | *MYO10* | 22KB | 0.06 |
| **chr5:16957447** | rs76494545 | A | 0.3 | 1.79E-7 | *MYO10* | 21KB | 0.06 |
| **chr5:16959472** | rs74669701 | A | 0.3 | 1.99E-7 | *MYO10* | 23KB | 0.06 |
| **chr5:16959654** | rs78658115 | C | 0.3 | 1.99E-7 | *MYO10* | 23KB | 0.06 |
| **chr5:16924422** | rs17651767 | G | 0.3 | 4.47E-7 | *MYO10* | Intron 1 | 0.06 |
| **chr5:16949802** | rs76015545 | C | 0.23 | 7.47E-6 | *MYO10* | 13KB | 0.07 |

TSS: transcription start site; AF: Allele frequency from gnomAD

### Integrated analysis suggests SNPs associate with regulatory regions for *MYO10*

As these variants were in a single intergenic association peak on chromosome 5 near *MYO10*, the LD structure around the variants was evaluated. Significant variants rs112072727 and rs78565023 form an LD block with variant rs74669701 (pink lettering) (**Figure 5A)**. A transcription factor binding site analysis reveals a putative SOX9 binding site is created at rs74669701 after the deletion of a T in the alternative sequence (**Figure 5B**, pink box). The closest intergenic variant was positioned 13KB 5’ of the *MYO10* transcriptional start site (**Table 3**). H3K27Ac data from human heart left ventricle and subcutaneous abdominal adipose tissue showed enrichment in and around intronic variant rs17651767 (**Figure 6A**, blue and black tracks, yellow box). Data from human aortae showed enrichment of H3K27Ac peaks in and around rs76494545 and rs113861938 (**Figure 6A**, green track, light blue box). H3K27Ac enrichment in and around rs76494545 and rs113861938 was also seen in data from thoracic, and ascending aorta. ATAC-seq data from iPSC-derived cardiomyocytes did not show activity in the regions of interest (gold and light blue boxes) indicating that iPSC-derived cardiomyocytes are likely not an appropriate model of *MYO10* gene regulation (**Figure 6A**, red track).

**Figure 5.**
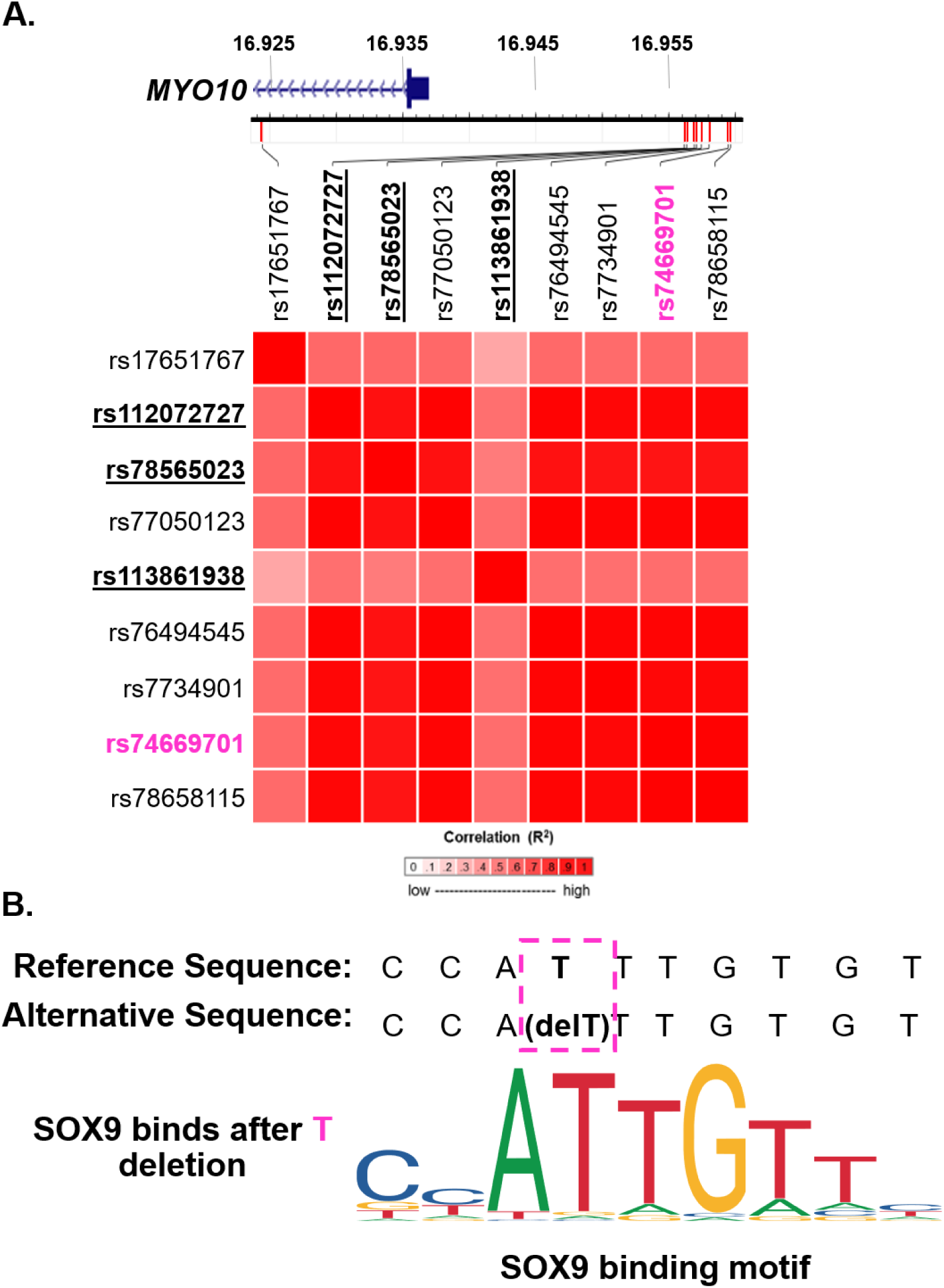
LD structure and location of SNPs associated with LVIDd and *MYO10*. **A.** The position of *MYO10* is shown. Shown are SNPs on chromosome 5 in LD with the SNPs significantly associated with LVID trajectory probability in all 1000 Genomes Populations (https://ldlink.nci.nih.gov/). SNPs that were significantly associated (p-value<3E-08) with LVIDd trajectory probability; rs112072727, rs78565023, rs113861938; are bolded and underlined. Variants rs112072727 and rs78565023 form an LD block with variant rs74669701 (pink lettering). **B.** A transcription factor binding site analysis showed that a putative SOX9 binding site is created at rs74669701 (pink lettering in **A** after the deletion of T in the alternative sequence (pink box).

**Figure 6.**
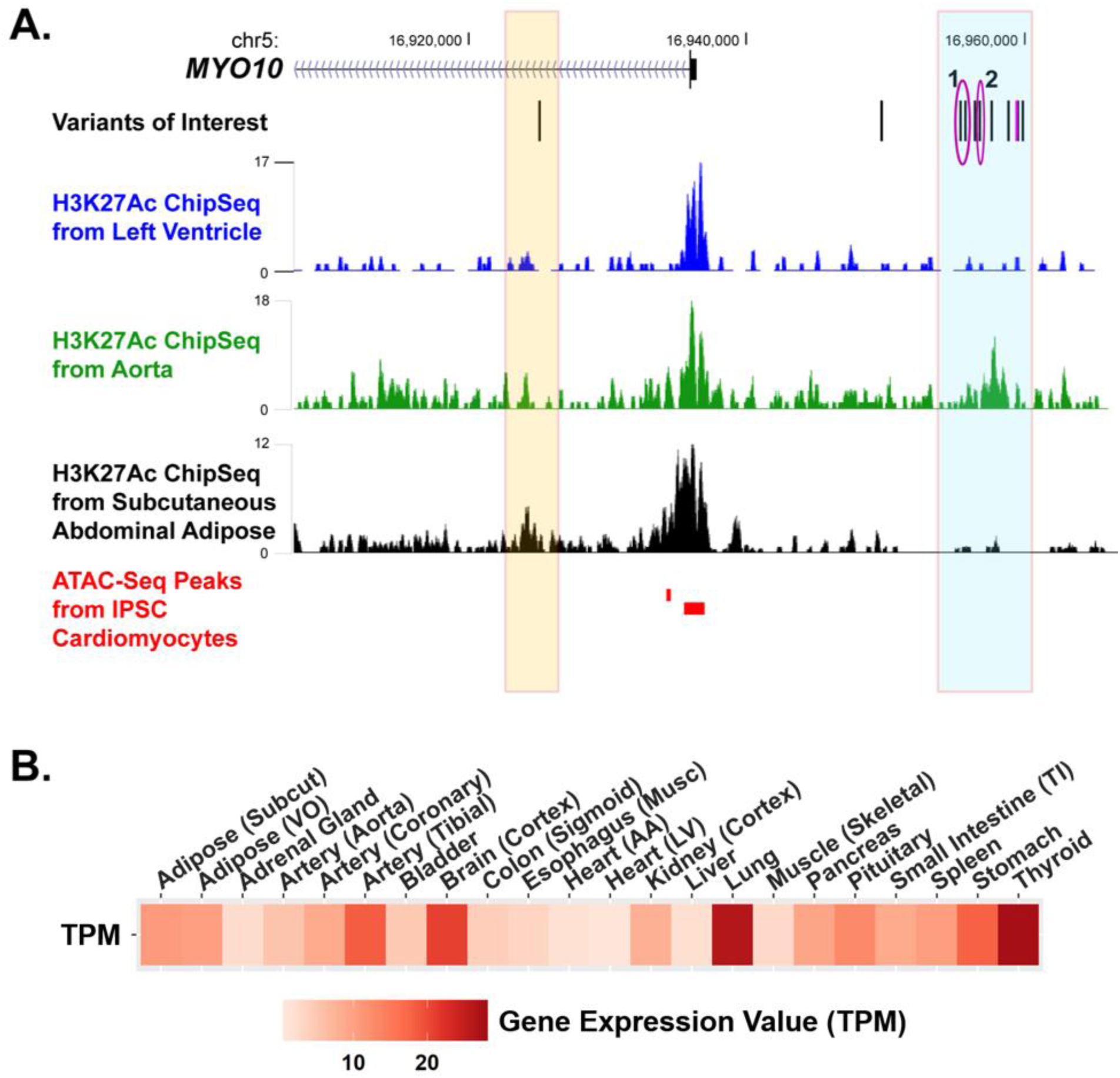
Localization of LVIDd-associated SNPs in putative *MYO10 e*nhancer regions. **A.** LVIDd-associated SNPs are denoted by black lines. Significant SNPs are circled in magenta. The gold and light blue boxes encompass putative enhancers that correspond with the SNPs found to be associated with LVIDd trajectory probability. Shown are H3K27Ac marks from human heart left ventricle in the blue track, from human aorta in the green track, and from human subcutaneous abdominal adipose in the black track. H3K27Ac data from the aorta showed enrichment for H3K27Ac marks in and around rs76494545 (magenta circle 1) and rs113861938 (magenta circle 2). H3K27Ac marks from left ventricle and subcutaneous abdominal adipose tissue show active enhancer peaks in and around variant rs17651767 (gold box). **B**. GTEx median expression in transcripts per million (TPM) of *MYO10* across all available tissue types (data from GTEx Portal on 3/28/2020). Abbreviations: Subcut-subcutaneous; VO-visceral(omentum); Musc-muscularis layer; AA-atrial appendage; LV-left ventricle; TI-terminal ileum

*MYO10* is ubiquitously expressed with high expression in thyroid, lung and tibial arteries (**Figure 6B**). *MYO10* is expressed in hearts at lower levels in the whole heart tissue. RNAseq data from 12 failed hearts and 3 healthy hearts, showed that *MYO10* was upregulated in failed hearts (log fold change=0.71, FDR adjusted p-value=0.038). The finding was validated in the larger cohort of 97 failed and 108 non-failed hearts (log fold change=0.086, Benjamini– Hochberg adjusted p-value=5.7E-11) (Heinig et al., 2017). An eQTL analysis using GTEx data show a significant association with *MYO10* and rs74669701, a variant in LD with variants significantly associated with LVIDd trajectory probability (**Figure 5A**), in the tibial artery (eQTL p-value=0.00099) but not left ventricle of the heart nor skeletal muscle. These analyses reveal that *MYO10* variants are associated with regulatory regions in vascuar tissues, expression changes in artery and is differentially regulated in heart failure indicating a potential role in cardiac extrinsic factors promoting heart failure.

## DISCUSSION

### Trajectory probability for GWAS

Electronic health record (EHR) data, in conjunction with genetic information, is a powerful tool to gain insight into novel and established disease mechanisms. Genetic studies relying on phenotype data derived from the EHR capture can readily capture clinical data in a cross-sectional and case/control manner. While useful, these data can be enhanced by analyzing progression of disease traits. Longitudinal data may provide novel genetic associations informing disease outcomes. Here we show that quantifying the probability of these cluster phenotypes provides a novel technique for which to find significant associations in small cohorts with rich longitudinal data. In this study, we used individual level longitudinal data to create clusters and trajectory probabilities used for cluster assignment. We analyzed QTc interval and LVIDd corrected for BSA using trajectory probability as a continuous trait and found this increased the likelihood to find associations compared to only using cluster in GWAS.

### WLS associates with EKG intervals

We found that QTc interval trajectory probability was associated with variants in putative enhancer regions linked to *WLS*, a gene encoding Wntless, a Wnt receptor that participates in Wnt’s secretion and recycling (Port and Basler, 2010). Deeper analyses suggested that these variants map to active enhancers in left ventricle, aorta, subcutaneous abdominal fat, and iPSC-derived cardiomyocytes. *WLS* is highly expressed in tissues with smooth muscle enrichment, including vascular smooth muscle as in the aorta. The high expression in aorta supports a vascularly-mediated mechanism to regulate cardiovascular outcomes, for example shifts in blood pressure may indirectly lead to myocardial compliance and fibrosis. The increase in *WLS* expression in failed hearts compared to non-failed hearts could reflect the non-cardiomyocyte component of these hearts including vasculature. *WLS* was previously associated with QT interval in a meta-analysis of Hispanic populations totaling approximately 16,000 individuals (Mendez-Giraldez et al., 2017). Specifically, the variant rs112611436, was associated with a 12ms increase in QT interval. In an animal model *WLS-* deficient macrophages were found to have enhanced anti-inflammatory and reparative properties in a mouse model of myocardial infarction (Palevski et al., 2017). Taken together, these data support an indirect mechanism by which *WLS* influences the myocardium and thereby the cardiac conduction system.

### MYO10 associates with increased LV dimensions

Trajectory probability GWAS using left ventricular dimensions (LVIDd) identified variants in putative enhancers for *MYO10*. A suggestive signal was found in the first intron of *MYO10*, a region that is known to be enriched for active transcriptional regulatory signals (Park et al., 2014). *MYO10* encodes a member of the myosin superfamily and the chromatin marks suggested *MYO10* expression in aorta and subcutaneous abdominal fat as the target tissues. *MYO10* expression was also increased in failed compared to non-failed hearts, which includes non-cardiomyocyte components. Variant rs74669701 showed an increase likelihood for binding of the SOX9 transcription factor which has been shown in previous literature to increase cardiac fibrosis in injured hearts (Lacraz et al., 2017; Scharf et al., 2019). The expression of *MYO10* in thyroid, lung and tibial arteries suggests a role as cardiac-extrinsic factor in promoting an enlarged heart. An intronic SNP in *MYO10* was previously associated with insulin-like growth factor binding protein 1 (IGFBP-1) ng/mL (Comuzzie et al., 2012). IGFBP-1 is a marker for insulin-resistance, and this study showed a suggestive association in a family-based cohort study of obesity in Hispanic children. Specifically, the variant, rs17614462, was associated with an increase in IGFBP-1, and this biomarker has been linked to increased risk of metabolic syndrome and cardiovascular diseases (Bae et al., 2013). Another intronic variant in M*YO10*, rs2434960, was previously associated with an increase in adiposity and leptin levels (Zhang et al., 2013). Metabolic syndrome is known to adversely affect HF outcomes and, therefore, these studies may provide insight into the genetic interplay between metabolic syndrome and HF (Perrone-Filardi et al., 2015). Metabolic syndrome has been previously shown to be associated with left ventricular septal thickness (Burchfiel et al., 2005), indicating a possible mechanism for which this gene is associated with cardiac outcomes.

### Conclusions and Study Limitations

Smaller cohorts can often be valuable because of having deeper phenotypic characterization. Longitudinal data can be especially powerful because it captures more of the data from electronic health records. Utilizing biobanks that are linked to EHRs provides a unique opportunity for phenotyping as there are a wealth of data that can be used in many ways especially longitudinally. Recent polygenic risk score (PRS) studies have shown that including longitudinal data in PRS increases the likelihood of finding associations (Zhao et al., 2019). Using longitudinal data in a method that is robust to missing data in a censored normal distribution using multivariable mixture model provides a novel method for which to detect genetic variants associated with cardiac traits. In addition, we found that using trajectory probabilities provides a method for which to find novel significant associations in small, well-phenotyped, longitudinal cohorts, such as those found in biobanks. This approach should prove useful in the study of rare disease cohorts and underrepresented populations.

One limitation of these analyses is that they only include the autosomes. As there are known sex differences in cardiovascular disease, it is possible that there are unexplored variants on the X chromosome that contribute to phenotype. We also only used the first three principal components to correct for global genetic ancestry. While these first 3 account for large substructure in the dataset, it is possible that adding additional components would further account for minor substructure in the data being analyzed. Lastly, unknown confounders could be contributing to the signal that are found in these association analyses.

## ACKNOWLEDGEMENTS

We thank the McDonnell Genome Institute at Washington University in St. Louis. We also gratefully acknowledge the participation of the NUgene biobank participants.

## FUNDING SOURCES

This work was supported by grants from the National Institutes of Health: NIH U01HG008673 (RC/MES), NIH R01HL128075 (EMM/MAN), NIH/NLM T32 LM012203 (TDP), NIH/NIDDK T32 DK007169 (TDP), and the American Heart Association 18CDA34110460 (MJP). The funders had no role in the study design, data collection and analysis, decision to publish, or preparation of the manuscript.

## DISCLOSURES

All participants provided written consent for participation in NUgene, and this work was performed under the ethical and regulatory approval of Northwestern University’s Institutional Review board (STU0010003).

## SUPPLEMENTAL INFORMATION FOR POTTINGER ET AL

**Supplemental Table 1.** ENCODE and Roadmap histone modification and chromatin organization datasets.

| Target | Dataset | Accession Number | Reference |
| --- | --- | --- | --- |
| H3K27Ac Histone Modification | Human LV- ChIP-Seq | GSM906396 | Roadmap Epigenomics Consortium, et al. 2015 |
| H3K27Ac Histone Modification | Human Aorta- ChIP-Seq | GSM906392 | Roadmap Epigenomics Consortium, et al. 2015 |
| H3K27Ac Histone Modification | Human Subcutaneous Abdominal Adipose- ChIP-Seq | ENCFF190GWA | ENCODE Project. 2012 |
| Open Chromatin | iPSC-CM- ATAC-Seq | E-MTAB-8983 | Montefiori L. & Nobrega |
| 3D Chromatin Organization | *iPSC-CM- Hi-C Promoter Capture* | E-MTAB-6014 | Montefiori L. et al 2018 |

**Supplemental Table 2.** GWAS SNPs associated with QTc interval trajectory probability, the corresponding associations in QTc interval cluster, and the first QTc interval measurement.

| Variant | rs number | Risk Allele | Parameter estimate QTc interval trajectory probability | P-value | Parameter estimate QTc interval cluster | P-value | Parameter estimate QTc interval measure 1 | P-value |
| --- | --- | --- | --- | --- | --- | --- | --- | --- |
| chr1:68762502 | rs1304851468 | G | 0.2 | 1.1E-8 | 3.8 | 1.5E-5 | 9.4 | 2.5E-3 |
| chr1:68788564 | rs4536035 | C | 0.19 | 1.8E-8 | 3.4 | 1.1E-5 | 8.5 | 3.3E-3 |
| chr1:68788712 | rs4506498 | T | 0.19 | 2.5E-8 | 3.4 | 1.3E-5 | 8.4 | 3.6E-3 |
| chr1:68785213 | rs11209261 | T | 0.18 | 6.3E-8 | 3.3 | 2E-5 | 8.2 | 4.8E-3 |
| chr1:68764949 | rs10889761 | T | 0.18 | 10E-8 | 3.2 | 4E-5 | 7.8 | 9.4E-3 |
| chr1:68797643 | rs10789260 | G | 0.19 | 1.8E-7 | 3.5 | 4E-5 | 8.8 | 4.9E-3 |
| chr1:68804349 | rs4926357 | G | 0.18 | 3E-7 | 3.3 | 6.6E-5 | 9.7 | 1.8E-3 |
| chr1:68787424 | rs11309689 | AT | 0.15 | 4.2E-7 | 2.9 | 3.9E-5 | 8 | 2.7E-3 |
| chr1:68784444 | rs6588315 | G | 0.17 | 4.3E-7 | 2.9 | 7.9E-5 | 7.6 | 8.4E-3 |
| chr1:68804055 | rs4926330 | A | 0.18 | 5.3E-7 | 3.1 | 1.2E-4 | 9.2 | 3.3E-3 |
| chr1:68794008 | rs4598529 | T | 0.18 | 5.5E-7 | 3.3 | 7.3E-5 | 8.7 | 5.9E-3 |
| chr1:68787891 | rs4244620 | C | 0.16 | 6.3E-7 | 3 | 5.1E-5 | 7.5 | 6.8E-3 |
| chr1:68804530 | rs4926360 | T | 0.18 | 7E-7 | 3.1 | 1.2E-4 | 9.3 | 2.7E-3 |
| chr1:68804258 | rs4926356 | A | 0.18 | 8.1E-7 | 3.1 | 1.3E-4 | 8.9 | 4.8E-3 |

**Supplemental Table 3.** GWAS SNPs associated with QTc interval trajectory probability by genetic ancestry.

| rs number | Risk Allele | Parameter estimate QTc interval trajectory probability (African) | P-value | Parameter estimate QTc interval trajectory probability (Hispanic) | P-value | Parameter estimate QTc interval trajectory probability (European) | P-value | Parameter estimate QTc interval trajectory probability (Other) | P-value |
| --- | --- | --- | --- | --- | --- | --- | --- | --- | --- |
| rs1304851468 | G | 0.22 | 2.3E-4 | 0.04 | 0.6 | 0.23 | 1.4E-3 | 0.31 | 4.1E-3 |
| rs4536035 | C | 0.17 | 6.7E-4 | 0.09 | 0.26 | 0.23 | 1.3E-3 | 0.26 | 8.5E-3 |
| rs4506498 | T | 0.17 | 8.4E-4 | 0.09 | 0.26 | 0.23 | 1.3E-3 | 0.26 | 8.5E-3 |
| rs11209261 | T | 0.17 | 1.5E-3 | 0.09 | 0.26 | 0.23 | 1.3E-3 | 0.26 | 1.2E-2 |
| rs10889761 | T | 0.19 | 9.5E-4 | 0.1 | 0.19 | 0.2 | 4.3E-3 | 0.26 | 1.2E-2 |
| rs10789260 | G | 0.19 | 2.1E-3 | 0.09 | 0.23 | 0.18 | 2.2E-2 | 0.32 | 2.2E-3 |
| rs4926357 | G | 0.18 | 2.1E-3 | 0.09 | 0.23 | 0.17 | 2.2E-2 | 0.32 | 2.2E-3 |
| rs11309689 | AT | 0.12 | 1.1E-2 | 0.09 | 0.21 | 0.2 | 2.6E-3 | 0.28 | 1.6E-3 |
| rs6588315 | G | 0.14 | 6E-3 | 0.09 | 0.26 | 0.23 | 1.3E-3 | 0.26 | 1.2E-2 |
| rs4926330 | A | 0.16 | 5.7E-3 | 0.1 | 0.16 | 0.17 | 2.2E-2 | 0.32 | 2.2E-3 |
| rs4598529 | T | 0.17 | 6.2E-3 | 0.09 | 0.23 | 0.18 | 2.2E-2 | 0.32 | 2.2E-3 |
| rs4244620 | C | 0.12 | 9.9E-3 | 0.09 | 0.26 | 0.23 | 1.3E-3 | 0.26 | 8.5E-3 |
| rs4926360 | T | 0.16 | 7.1E-3 | 0.09 | 0.23 | 0.18 | 2.2E-2 | 0.33 | 3.3E-3 |
| rs4926356 | A | 0.16 | 5.7E-3 | 0.09 | 0.23 | 0.17 | 2.2E-2 | 0.32 | 2.2E-3 |

**Supplemental Table 4.** GWAS SNPs associated with LVIDd trajectory probability, the corresponding associations in LVIDd cluster, and the first LVIDd measurement

| Variant | rs number | Risk Allele | Parameter estimate LVIDd trajectory probability | P-value | Parameter estimate LVIDd cluster | P-value | Parameter estimate LVIDd measure 1 | P-value |
| --- | --- | --- | --- | --- | --- | --- | --- | --- |
| **chr5:16957078** | rs113861938 | T | 0.42 | 2.1E-9 | 9.18 | 6.3E-6 | 0.4 | 4E-7 |
| **chr5:16956160** | rs112072727 | T | 0.33 | 1.3E-8 | 6.34 | 4.5E-6 | 0.32 | 5.8E-7 |
| **chr5:16956338** | rs78565023 | G | 0.32 | 1.4E-8 | 6.2 | 5.2E-6 | 0.32 | 5E-7 |
| **chr5:16956916** | rs77050123 | A | 0.31 | 6.5E-8 | 5.63 | 1.5E-5 | 0.31 | 2.1E-6 |
| **chr5:16958033** | rs7734901 | G | 0.3 | 7.5E-8 | 5.6 | 1.5E-5 | 0.29 | 4.4E-6 |
| **chr5:16957447** | rs76494545 | A | 0.3 | 1.8E-7 | 5.12 | 4.3E-5 | 0.3 | 5E-6 |
| **chr5:16959472** | rs74669701 | A | 0.3 | 2E-7 | 5.1 | 4.4E-5 | 0.28 | 9.9E-6 |
| **chr5:16959654** | rs78658115 | C | 0.3 | 2E-7 | 5.1 | 4.4E-5 | 0.28 | 9.9E-6 |
| **chr5:16924422** | rs17651767 | G | 0.3 | 4.5E-7 | 4.71 | 6.8E-5 | 0.27 | 5E-5 |
| **chr5:16949802** | rs76015545 | C | 0.23 | 7.5E-6 | 3.87 | 1.5E-4 | 0.26 | 9E-6 |

All values of LVIDd are corrected for BSA

**Supplemental Table 5.** GWAS SNPs associated with LVIDd trajectory probability by genetic ancestry.

| rs number | Risk Allele | Parameter estimate LVIDd trajectory probability (African) | P-value | Parameter estimate LVIDd trajectory probability (European) | P-value | Parameter estimate LVIDd trajectory probability (Hispanic) | P-value | Parameter estimate LVIDd trajectory probability (Other) | P-value |
| --- | --- | --- | --- | --- | --- | --- | --- | --- | --- |
| rs113861938 | T | 0.61 | 1.1E-6 | 0.34 | 4.9E-3 | 0.17 | 0.3 | 0.66 | 2.2E-2 |
| rs112072727 | T | 0.47 | 5.2E-5 | 0.33 | 6.8E-3 | 0.15 | 0.1 | 0.6 | 7E-3 |
| rs78565023 | G | 0.47 | 4.8E-5 | 0.34 | 4.9E-3 | 0.13 | 0.2 | 0.6 | 7E-3 |
| rs77050123 | A | 0.47 | 4.8E-5 | 0.34 | 4.9E-3 | 0.1 | 0.3 | 0.6 | 7E-3 |
| rs7734901 | G | 0.47 | 4.8E-5 | 0.34 | 4.9E-3 | 0.1 | 0.3 | 0.43 | 1.2E-2 |
| rs76494545 | A | 0.47 | 4.8E-5 | 0.34 | 4.9E-3 | 0.1 | 0.3 | 0.66 | 2.2E-2 |
| rs74669701 | A | 0.47 | 4.8E-5 | 0.34 | 4.9E-3 | 0.1 | 0.3 | 0.42 | 3.6E-2 |
| rs78658115 | C | 0.47 | 4.8E-5 | 0.34 | 4.9E-3 | 0.1 | 0.3 | 0.42 | 3.6E-2 |
| rs17651767 | G | 0.32 | 1.9E-3 | 0.28 | 1.6E-2 | 0.15 | 0.17 | 0.6 | 7E-3 |
| rs76015545 | C | 0.11 | 0.1 | 0.39 | 4.2E-3 | 0.12 | 0.17 | 0.51 | 1.2E-2 |

All values of LVIDd are corrected for BSA

## SUPPLEMENTAL FIGURES

**Supplemental Figure 1.**
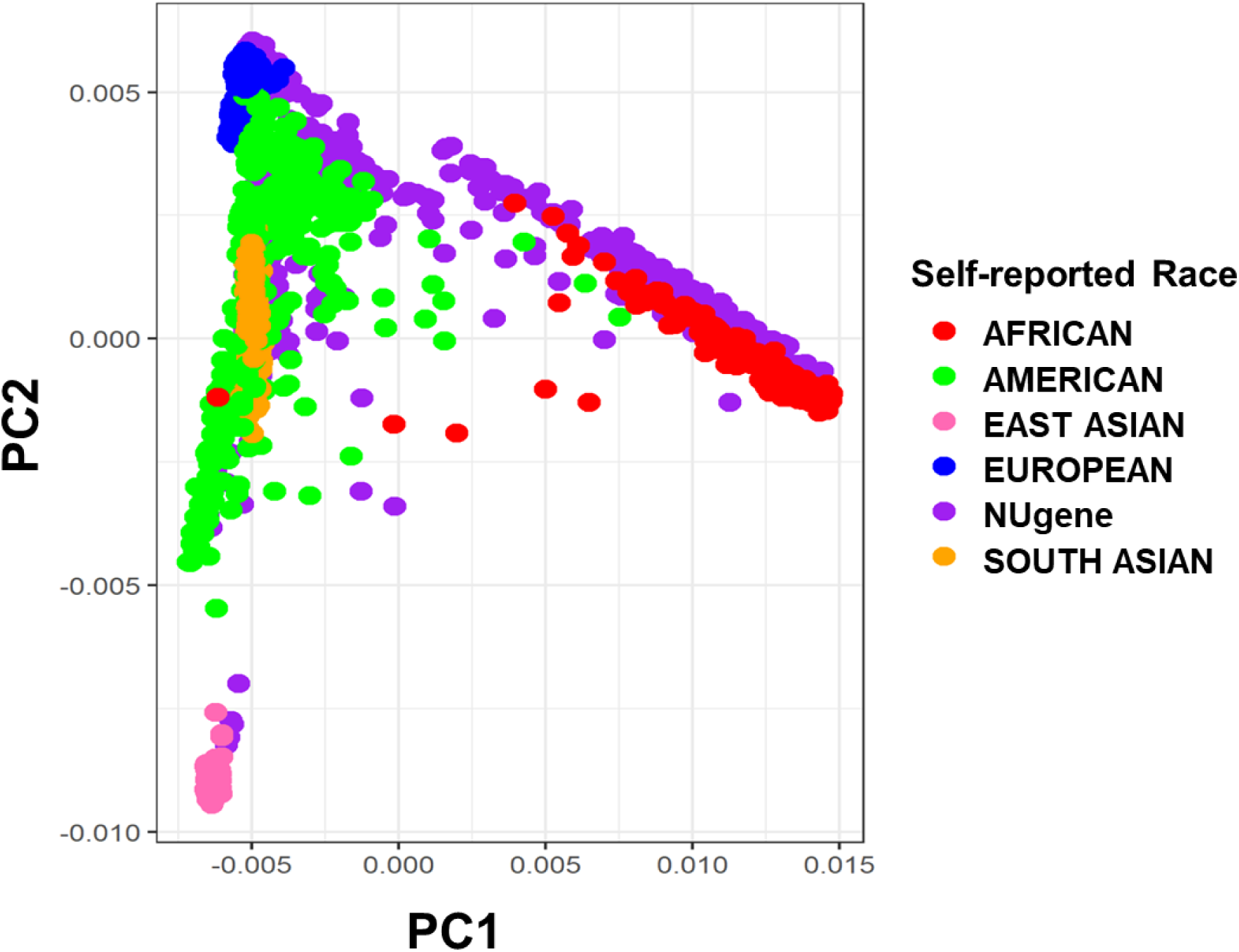
The NUgene cohort was anchored to the 1000 Genomes data and a principal components analysis was performed. This analysis shows that the NUgene cohort aligns well with the 1000 Genomes data. The individuals in the NUgene Cohort who cluster with the 1000 Genomes African cluster showed more European admixture than those identified in 1000 Genomes data.

**Supplemental Figure 2.**
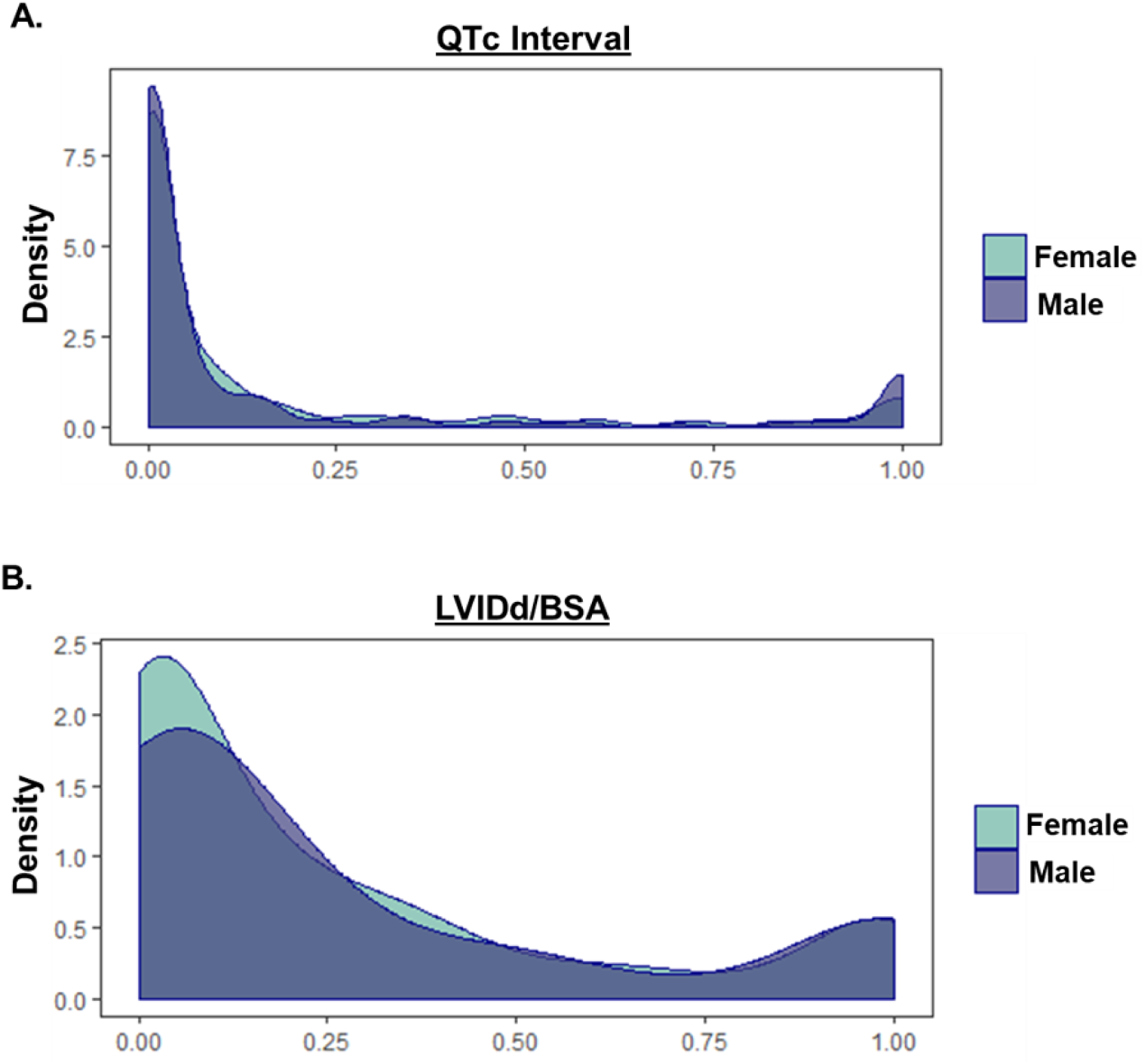
QTc Interval phenotyping and LVIDd/BSA phenotyping sex differences. **A.** The probability estimate used to group individuals with measures of QTc Interval into 2 groups, normal and long QTc interval, show a distribution between 0 and 1. There are no statistical differences in trajectory probabilities between males in females in this study. **B.** The probability estimate used to group individuals with measures of LVIDd/BSA into 2 groups, normal and dilated LVIDd/BSA, show a distribution between 0 and 1. There are no statistical differences in trajectory probabilities between males in females in this study.

**Supplemental Figure 3.**
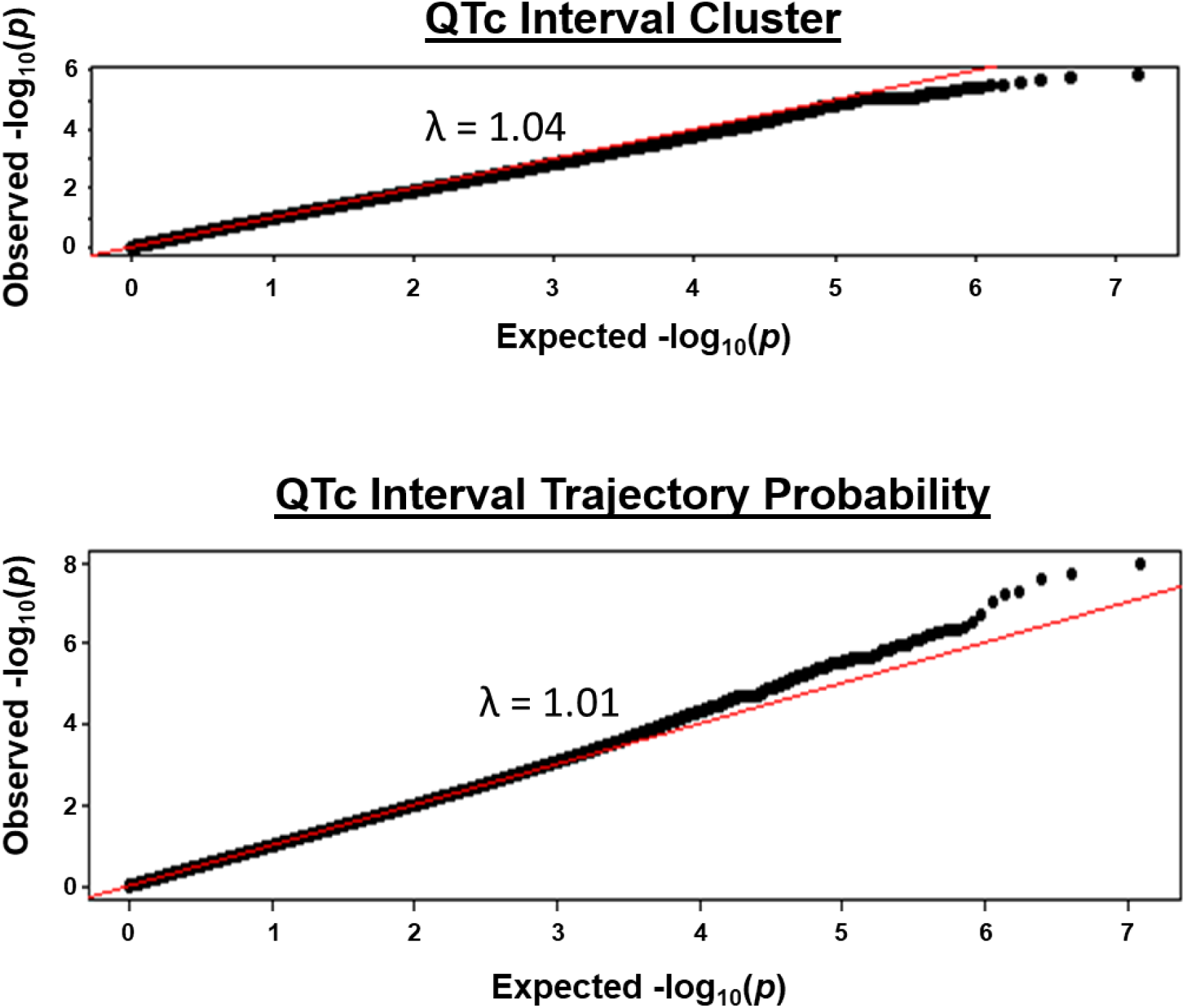
Quantile-Quantile plot of the GWAS of QTc interval cluster and QTc interval trajectory probability. **A.** QQ-plot of the GWAS of QTc interval cluster shows slight deviations from normal (λ = 1.04). **B.** QQ-plot of the GWAS of QTc interval trajectory probability shows minor deviations from normal (λ = 1.01).

**Supplemental Figure 4.**
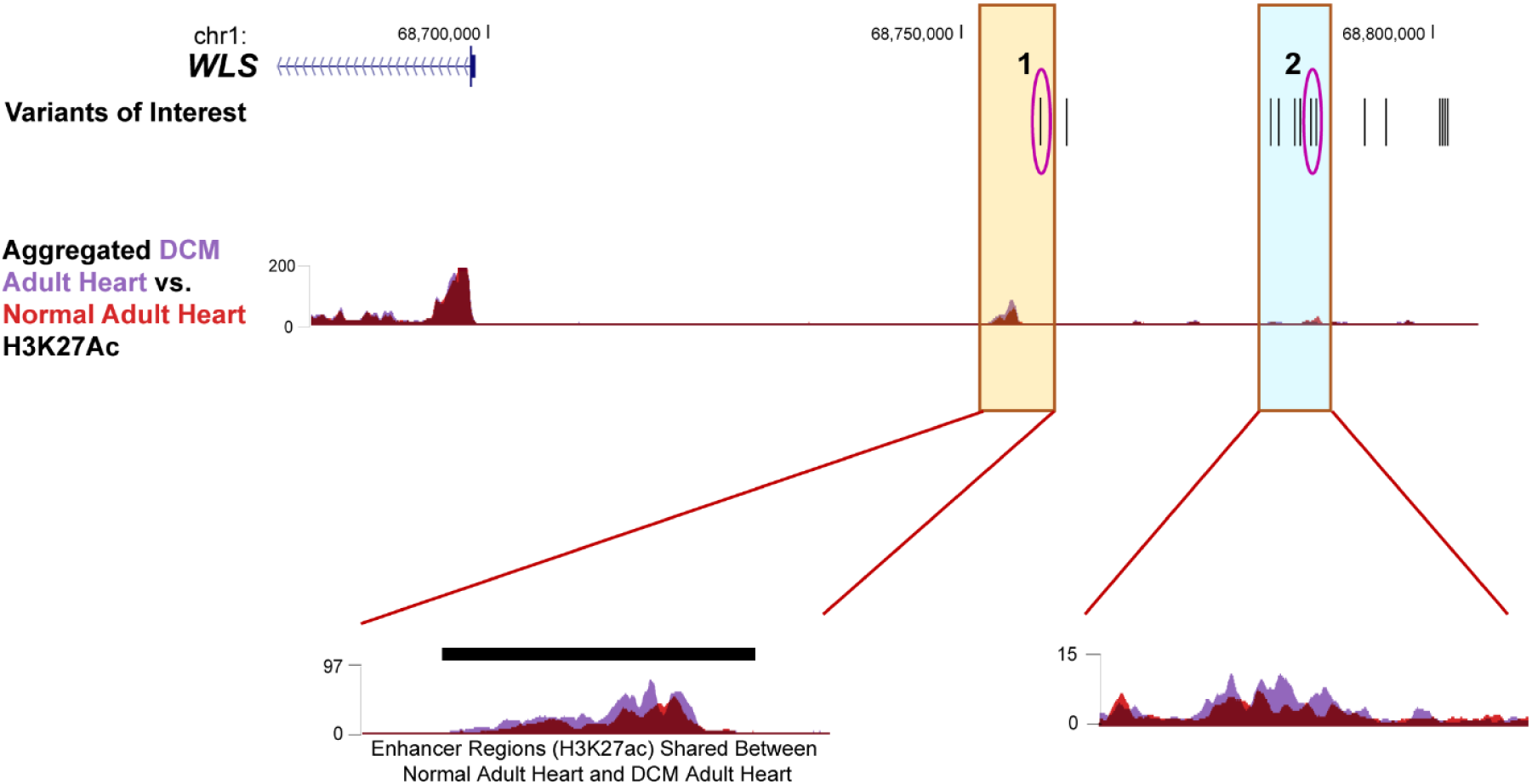
Localization of QTc interval-associated SNPs in putative *WLS* enhancer regions in normal human adult hearts and hearts with DCM. H3K27Ac Chip-Seq data in human adult non-failed hearts (red) and failed hearts (light purple) showed the acetylation levels of putative regulatory regions between the different heart samples. Insets show putative enhancer regions (gold and light blue boxes) which reveal that non-failed and failed hearts have similar acetylation patterns in the proximal putative regulatory region (gold box, black bar), while the pattern in the distal region (blue box) was neither significantly similar or different per this analysis (Spurrell, et al. 2019).

**Supplemental Figure 5.**
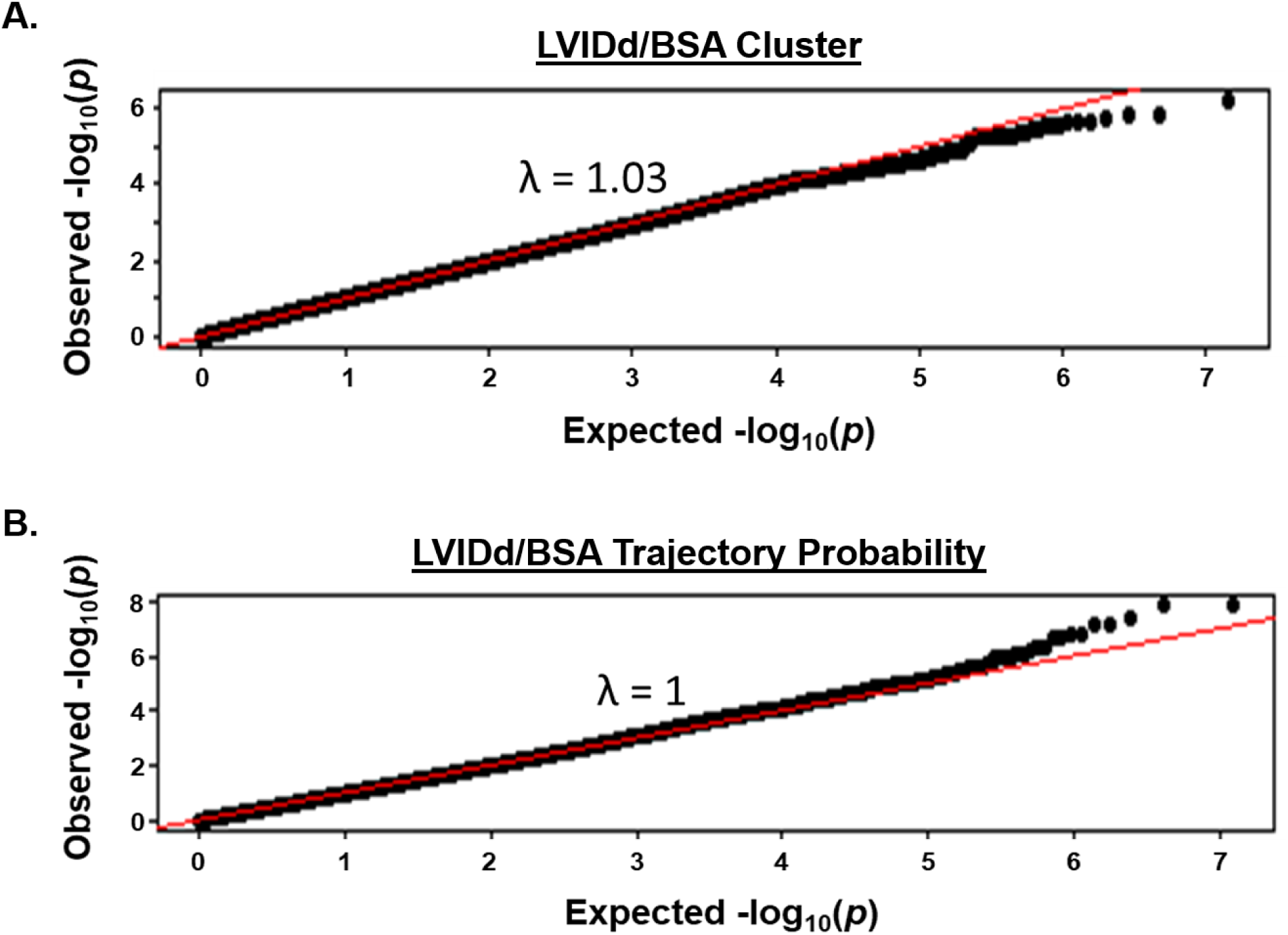
Quantile-Quantile plot of the GWAS of LVIDd/BSA cluster and LVIDd/BSA trajectory probability. **A.** QQ-plot of the GWAS of LVIDd/BSA cluster shows slight deviations from normal (λ = 1.03). **B.** QQ-plot of the GWAS of LVIDd/BSA trajectory probability shows no deviation from normal (λ = 1).

## Notes

### Competing Interest Statement

The authors have declared no competing interest.

